# Sarcolambans are phospholamban- and sarcolipin-like regulators of the sarcoplasmic reticulum calcium pump SERCA

**DOI:** 10.1101/2020.07.19.211243

**Authors:** Jessi J. Bak, Rodrigo Aguayo-Ortiz, Muhammad Bashir Khan, Seth L. Robia, M. Joanne Lemieux, L. Michel Espinoza-Fonseca, Howard S. Young

## Abstract

From insects to humans, calcium signaling is essential for life. An important part of this process is the sarco-endoplasmic reticulum calcium pump SERCA, which maintains low cytosolic calcium levels required for intracellular calcium homeostasis. In higher organisms, this is a tightly controlled system where SERCA interacts with tissuespecific regulatory subunits such as phospholamban in cardiac muscle and sarcolipin in skeletal muscle. With the recent discovery of the sarcolambans, the family of calcium pump regulatory subunits also appears to be ancient, spanning more than 550 million years of evolutionary divergence from insects to humans. This evolutionary divergence is reflected in the peptide sequences, which vary enormously from one another and range from vaguely phospholamban-like to vaguely sarcolipin-like. Here, our goal was to investigate select sarcolamban peptides for their ability to regulate calcium pump activity. For a side-by-side comparison of diverse sarcolamban peptides, we tested them against mammalian skeletal muscle SERCA1a. This allowed us to determine if the sarcolamban peptides mimic phospholamban and sarcolipin in their regulatory activities. Four sarcolamban peptides were chosen from different invertebrate species. Of these, we were able to express and purify sarcolamban peptides from bumble bee, water flea, and tadpole shrimp. Sarcolamban peptides were co-reconstituted into proteoliposomes with mammalian SERCA1a and the effect of each peptide on the apparent calcium affinity and maximal activity of SERCA was measured. While all peptides were super-inhibitors of SERCA, they exhibited either phospholamban-like or sarcolipin-like characteristics. Molecular modeling, protein-protein docking, and molecular dynamics simulations were used to reveal novel features of insect versus mammalian calcium pumps and the sarcolamban regulatory subunits.

## INTRODUCTION

Sarcolambans (SLB) are a group of membrane peptides conserved from insects to humans and encoded by small open-reading frames (smORF) in what were thought to be noncoding RNAs (1). Recently, several groups have established that some noncoding RNAs are misannotated and contain smORFs that encode biologically active peptides. The first such peptides were found to play a role in *Drosophila* development (2–4), though the wider significance of smORFs and small peptides across species remained an open question. The discovery of SLBs and their implication in muscle function answered this question on the conservation of smORFs and the peptides they encode (1). Since these initial findings, so-called micropeptides have been found to play important roles in a variety of biological processes, leading to the speculation that they are much more common than currently understood (5–7). However, the characterization of peptides and smORFs is challenging because of the abundance in plant and animal genomes and the difficulty in validating the translation and function of small peptides. While small functional peptides are known throughout nature, most of these were discovered by biochemical means and the encoding genomic DNA sequences were later found based on the peptide amino acid sequences. Excellent examples of this “grind and find” approach include phospholamban (PLN (8)) in cardiac muscle and sarcolipin (SLN (9)) in skeletal muscle. The further identification of smORFs and small peptides – less than 300 base pairs and 100 amino acids, respectively – has required the development of new bioinformatic and transcriptomic approaches to identify RNAs and validate the smORFs within. With these advances, the SLBs have been joined by additional membrane peptides hidden in “noncoding” RNAs, which regulate calcium transport in the sarco-endoplasmic reticulum (5–7). This may be a general physiological mechanism that is repeated throughout membrane transport biology. Indeed, another family of single span transmembrane peptides, the FXYD proteins, is important for modulating sodium and potassium transport across the plasma membrane (10–12)

SLB peptides are considered the invertebrate orthologues of the mammalian regulatory peptides PLN and SLN, with putative sequence and functional similarities to their mammalian counterparts. Bioinformatic analysis of putative noncoding RNAs in *Drosophila* revealed two potentially functional smORFs in a transcript expressed in somatic muscle and the post-embryonic heart (1). These peptides had the hydrophobicity and predicted helical characteristics of a transmembrane peptide, and they localized to the sarco-endoplasmic reticulum near dyad junctions. A combination of sequence analysis and structural homology showed that human sarcolipin (SLN), a regulator of SR calcium homeostasis in skeletal and atrial muscle, was a match to SLBs. The subsequent generation of sequence alignments and a phylogenetic tree revealed that SLBs, SLN, and the longer mammalian peptide, PLN, all originated from a single gene, making this study the first to show the conservation and evolution of a smORF. With this and the knowledge that SLN and PLN function to regulate calcium transport by the SR calcium pump SERCA, the SLBs were shown to be part of an ancient family of peptides that regulate calcium movement and whose function is required for normal muscle contraction in insects. The target of these peptides is the insect equivalent of SERCA, Ca-P60A, which is approximately 70% identical to mammalian SERCAs and has the same essential residues responsible for calcium binding and peptide regulation. The SLB peptides co-localize with *Drosophila* Ca-P60A and co-immunoprecipitate with it, and PLN can partially rescue the cardiac phenotype of a SLB knockout strain. Thus, SLBs and Ca-P60A appear to be the insect equivalents of PLN, SLN, and SERCA.

PLN and SLN are practically invariant across mammalian species, limiting our ability to use sequence variation as a guide for mutagenesis and structure-function analyses. The SLBs, however, span more than 550 million years of evolutionary divergence, and the known peptide sequences vary enormously from one another and range from more PLN-like to more SLN-like sequences. In *Drosophila*, two SLB peptides were found to regulate calcium and muscle contraction in the heart (1), much like PLN and SLN. With this in mind, our goal was to investigate how different SLB peptides alter the calcium-dependent ATPase activity of SERCA. Depending on how the SLB peptides alter the apparent calcium affinity and maximal activity of SERCA, in comparison to PLN and SLN, this would shed light on SLBs as calcium-transport regulators and identify structural elements that are important for their regulatory properties. To examine these regulatory properties and sequence variations, four different SLB peptides were chosen from different invertebrate species. Of these, we were able to express and purify SLB peptides from *Bombus terrestris* (bumble bee), *Daphnia pulex* (water flea), and *Triops cancriformis* (tadpole shrimp). Purified peptides, lipids, and SERCA were co-reconstituted into proteoliposomes and the effect of each peptide on the apparent calcium affinity and maximal activity of SERCA was measured. In doing this, it was found that each SLB peptide had a unique effect on SERCA, exhibiting either PLN-like or SLN-like characteristics. These functional differences are explained by sequence variations in the SLB peptides. More broadly, this work shows how the function of these peptides and the regulation of the sarcoplasmic reticulum calcium pump (Ca-P60A in insects (13)) is conserved across species.

## RESULTS

### Co-Reconstitution of SERCA and SLB Peptides

Four insect SLB orthologues were selected to study how sequence variation alters regulation of SERCA activity. We were unable to express and purify *Drosophila melanogaster* (fruit fly) SLB, though we readily obtained SLB peptides from *Bombus terrestris* (bumble bee), *Daphnia pulex* (water flea), and *Triops cancriformis* (tadpole shrimp) (**Figure 1**). SERCA was co-reconstituted into proteoliposomes in the absence and presence of the SLB peptides. The co-reconstitution procedure has been described (e.g. see (14–16)). The final molar ratio in the proteoliposomes was 1 SERCA to 120 lipids, and we targeted a 1:5 ratio of SERCA to SLB. The SLB peptides readily reconstituted with SERCA and subsequent SDS-PAGE revealed that the SLB peptides formed monomeric and oligomeric species (**Figure 2**). The efficiency of co-reconstitution was comparable to our previous studies with PLN (14,16,17) and SLN (15,18), though the level of Coomassie-staining varied somewhat between SLB peptides. Nonetheless, it was apparent that the SLB peptides co-reconstituted with SERCA and the resultant proteoliposomes were suitable for comparative functional studies with PLN and SLN.

**Figure 1:**
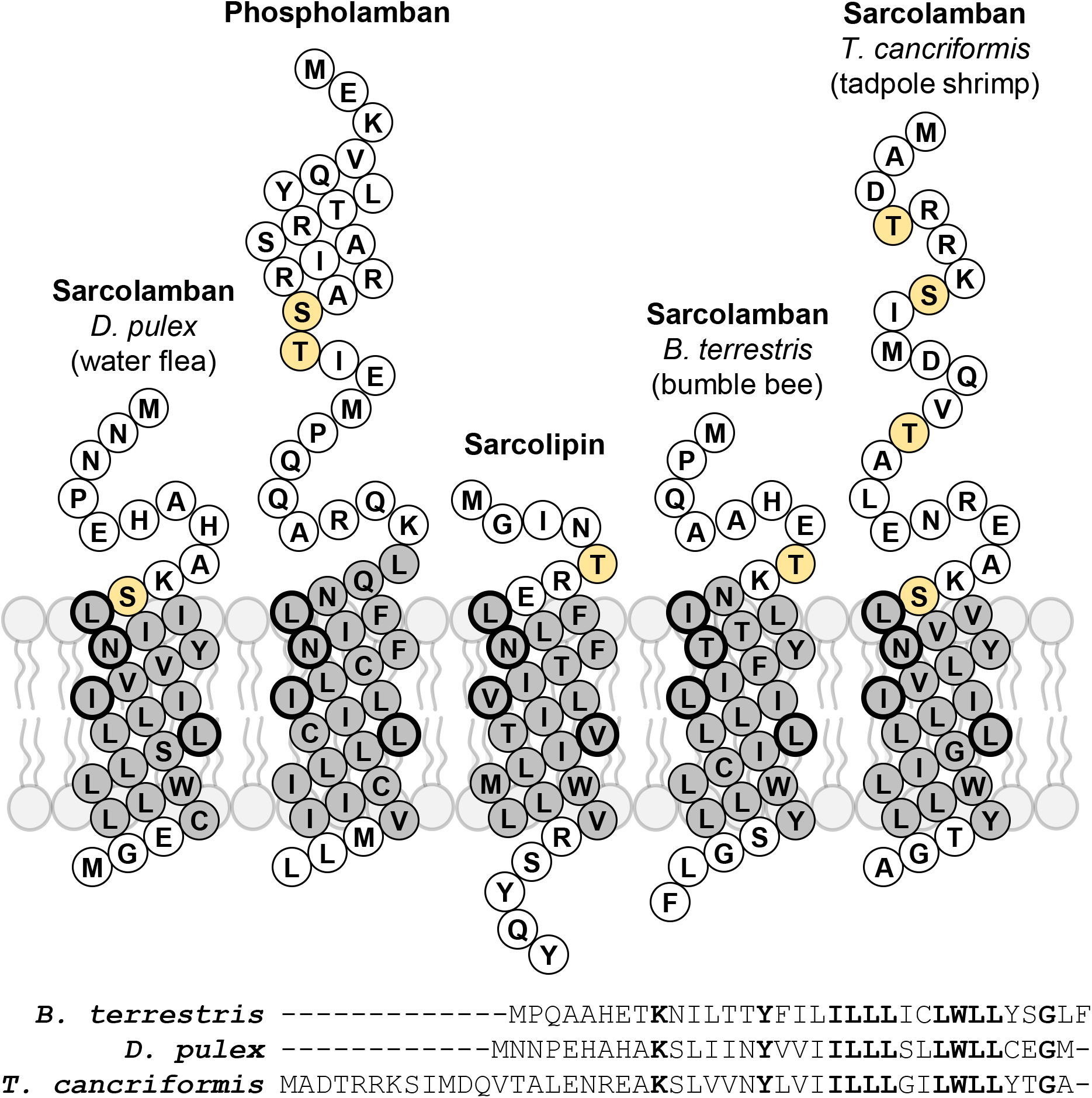
Topology diagrams for phospholamban (PLN), sarcolipin (SLN), and the sarcolamban (SLB) peptides used in the present study. A sequence alignment of the SLB peptides is also shown. The transmembrane domains are colored grey. The four essential residues in PLN are circled. When mutated to alanine, these residues have the largest impact on PLN function. Potential sites of regulatory control via phosphorylation are in orange.

With these proteoliposomes, the calcium-dependent ATPase activity of SERCA was measured, where SERCA reconstituted in the absence of peptides served as a negative control (**Figure 2**). Under these conditions, proteoliposomes containing SERCA alone yielded an apparent calcium affinity (K_Ca_) of 0.44 ± 0.03 μM calcium and a maximal activity (V_max_) of 4.1 ± 0.1 μmol/mg/min. Incorporation of bumble bee SLB into proteoliposomes with SERCA decreased both the apparent calcium affinity (K_Ca_ of 1.82 ± 0.15 μM calcium) and maximal activity of SERCA (1.1 ± 0.1 μmol/mg/min). Incorporation of water flea SLB into proteoliposomes with SERCA decreased the apparent calcium affinity (K_Ca_ of 1.53 ± 0.15 μM calcium) and increased the maximal activity of SERCA (V_max_ of 4.8 ± 0.2 μmol/mg/min). Finally, incorporation of tadpole shrimp SLB into proteoliposomes with SERCA decreased both the apparent calcium affinity (K_Ca_ of 2.05 ± 0.12 μM calcium) and maximal activity of SERCA (V_max_ of 2.3 ± 0.1 μmol/mg/min). These data are summarized in **Figure 3** and **Table 1**. Despite the sequence variation in the SLB peptides, they share a common feature of being super-inhibitors of SERCA. They differ in their abilities to regulate the maximal activity of SERCA, with bumble bee and tadpole shrimp SLB decreasing V_max_ and water flea SLB increasing V_max_.

**Figure 2:**
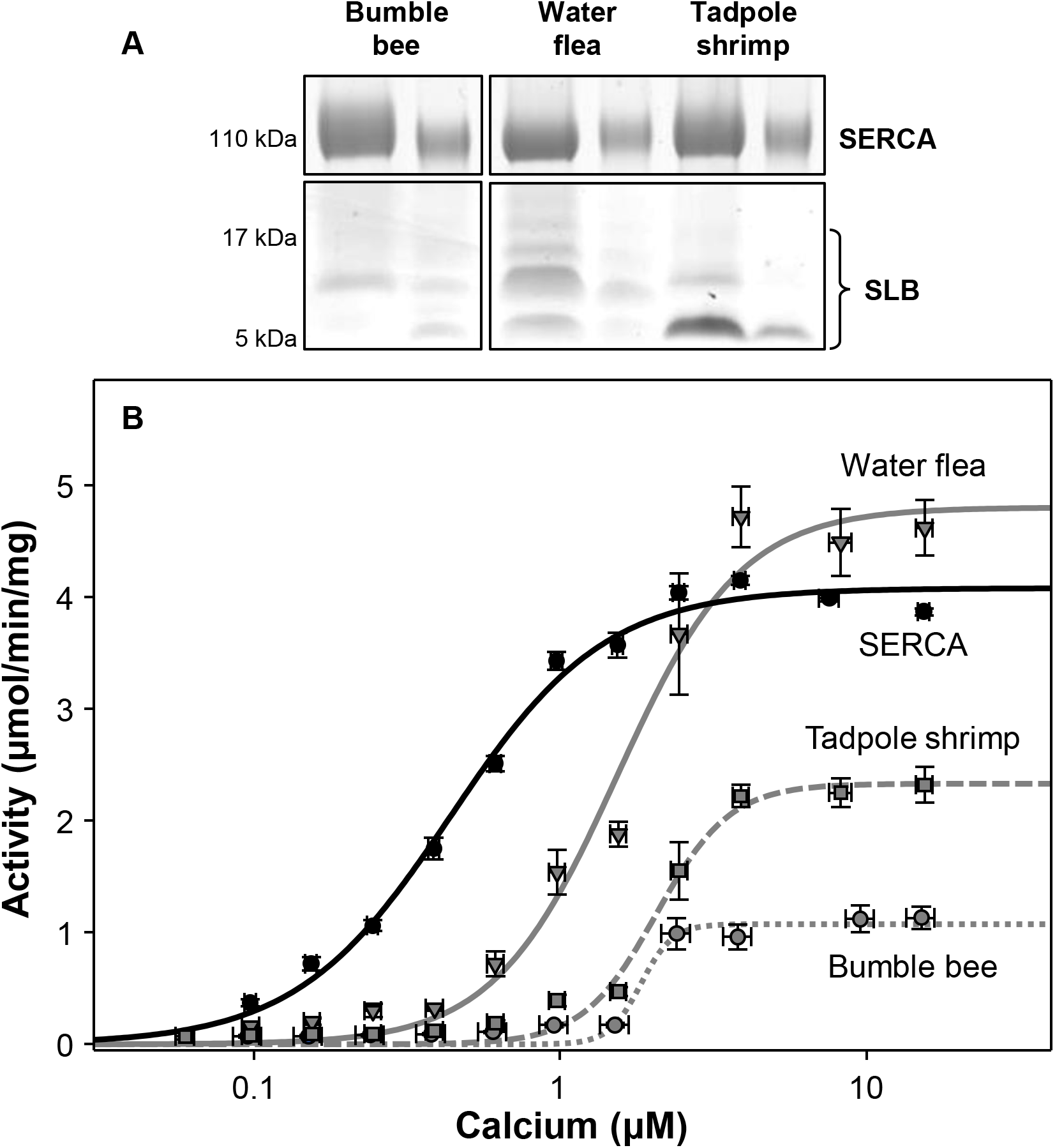
ATPase activity of co-reconstituted proteoliposomes containing SERCA and SLB peptides. (**A**) SDS-PAGE analysis of the proteoliposomes at two different protein concentrations. To visualize SERCA, a 10% polyacrylamide gel was run. To visualize peptides, a 15% polyacrylamide gel was run. (**B**) ATPase activity of SERCA reconstituted in the absence (SERCA; *black circles)* and in the presence of bumble bee SLB *(grey circles),* tadpole shrimp SLB *(grey squares’),* and water flea SLB *(grey triangles’).* Data points represent a minimum of three independent reconstitutions and a minimum of six ATPase activity assays (mean ± SEM). From these data, the calcium concentration at half-maximal activity (K_Ca_) and the maximal activity (V_max_) were calculated based on non-linear leastsquares fitting of the activity data to the Hill equation (**Table 1**).

**Table 1:**
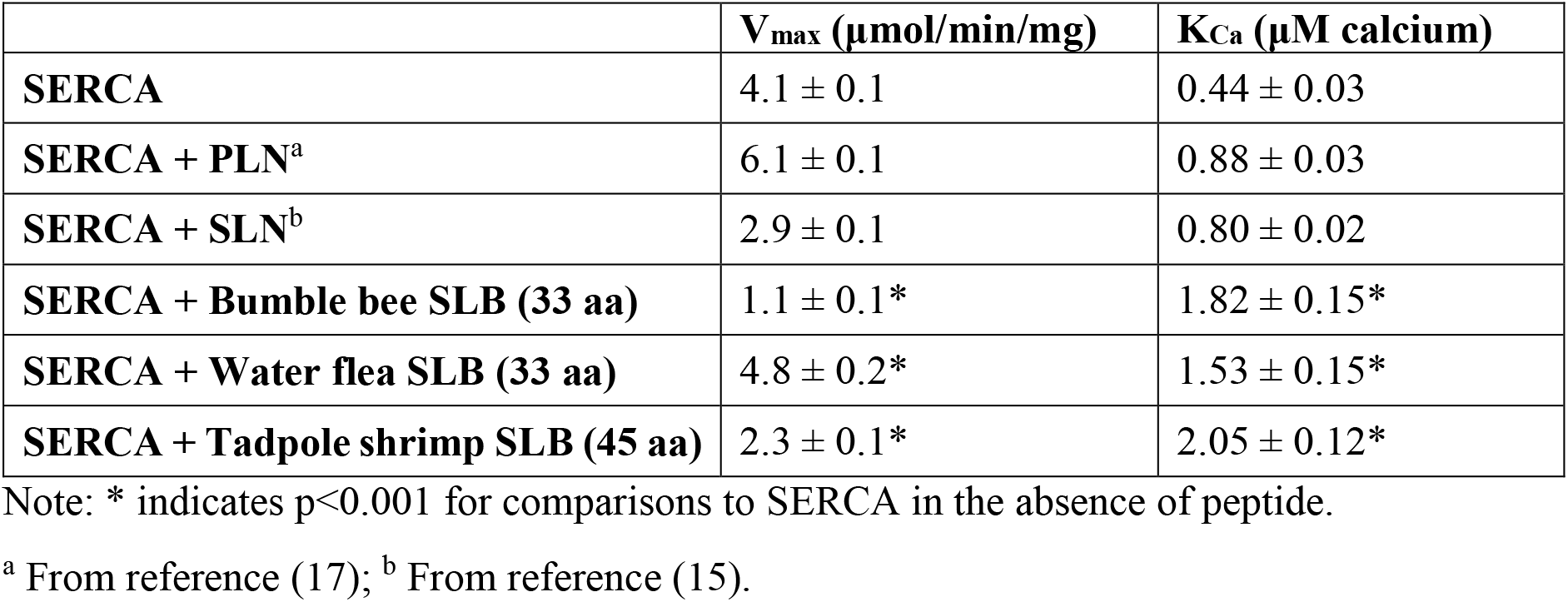
Kinetic parameters for SERCA reconstituted in the absence and presence of SLB peptides.

The three SLBs vary in length from 33 to 45 amino acids and they have 11 residues in common (**Figure 1**). Interesting features of the SLB peptides include a lysine residue at the N-terminus of the transmembrane domain (Lys^9^ in bumble bee SLB), which corresponds to Gln^29^ in PLN and Arg^6^ in SLN. In addition, the tryptophan found in SLN (Trp^23^) is conserved in the SLB peptides (Trp^26^ in bumble bee SLB). Finally, a series of conserved hydrophobic residues surround the tryptophan and make up the C-terminal half of the transmembrane domain (between residues 19-28 in bumble bee SLB). These structural elements are likely responsible for the observed super-inhibition of SERCA; however, these SLB peptides have different effects on the maximal activity of SERCA. Two regulators, bumble bee and tadpole shrimp SLB, were observed to be more SLN-like (they decrease V_max_) and one regulator, water flea, was observed to be more PLN-like (it increases V_max_). These observations are consistent with the sequence alignments, where water flea SLB is more homologous to PLN (**Figure 4**).

### Molecular modeling of Drosophila melanogaster Ca-P60A

To understand the super-inhibitory properties of the SLB peptides, we undertook molecular modeling on an insect SERCA, Ca-P60A, as well as protein-protein docking and molecular dynamics simulations of SERCA1a in an inhibitory complex with a SLB peptide. A molecular model of *D. melanogaster* calcium pump, Ca-P60A, was constructed using the Swiss-Model server (19) and rabbit SERCA1a in the E2 state as a template. The purpose of this exercise was to compare the binding groove for PLN and SLN between mammalian SERCA and the insect calcium pump Ca-P60A. The E2 state of SERCA was chosen as a model template because the binding groove for PLN and SLN is fully open. These calcium pump proteins are ~70% identical and the non-conserved residues are largely found on the surfaces of the protein (**Figure 5**). Indeed, the inhibitory binding groove (20–22) and the accessory binding site (23–25) for PLN and SLN are practically invariant between insects and mammals. In the inhibitory groove, sequence variations in Ca-P60A include Val^952^, Ser^954^, and Thr^955^ at the base of M9, which correspond to Pro^952^, Pro^954^, and Met^955^ in rabbit SERCA1a. These changes, particularly the proline residues in SERCA1a, appear to be responsible for the increased inhibitory capacity of the SLB peptides. At the accessory site, there is a single sequence variation Ala^273^, which corresponds to Leu^273^ in rabbit SERCA1a. Leu^273^ was previously implicated in the interaction of PLN and SLN with the M3 accessory site of SERCA (23,24).

### Molecular model and molecular dynamics simulations of a SERCA-SLB complex

Bumble bee SLB was used for molecular modeling and molecular dynamics simulations because it was the most inhibitory peptide. A structure model of bumble bee SLB peptide was built based on SLN in the crystal structures with SERCA (21,22), followed by alignment of the SLB model to SLN to generate a SERCA-SLB complex. The model was inserted in a lipid bilayer containing a total of 420 lipid molecules, the system was solvated, and potassium and sodium ions were added to reach a concentration of 150 mM and neutralize the total charge of the system. Molecular dynamics (MD) simulations of SERCA-SLB were carried out as described previously (23,24), and three independent 500 ns MD replicates of the SERCA-SLB complex were performed. The backbone RMSD for the SERCA-SLB complexes was stable for the last 300 ns of each simulation and the most representative structures of the complex were selected (**Figure 6A, B**). The conformation of SERCA changed in all three MD replicates (**Figure 6C**). Due to the lack of an ATP analog in the simulation, the N domain initially has a high solvent accessible surface area, which becomes stabilized by an interaction with the A domain. The residues involved include charge clusters on the A and N domains, including Arg^134^, Lys^135^, and Arg^139^ of the A domain and Asp^426^, Glu^429^, and Glu^466^ of the N domain. This interaction between the N and A domains causes the two domains to tilt toward the membrane. A similar domain movement occurs during the transition from E1 to E2 state of SERCA; however, in the simulation, the A domain does not rotate as observed for the E2 state.

Despite the cytoplasmic domain movements, the SERCA-SLB complex closely resembles the known SERCA-SLN complexes (21,22) Residues of SERCA and SLB that were within 3.0 Å of each other were determined from these merged MD trajectories and the representative structures of the SERCA-SLB complex (**Figure 6D, E**). Perhaps not surprising, the interaction of SLB with SERCA is very similar to SLN. The inclination of SLB is more upright and perpendicular to the membrane plane, while SLN is slightly more inclined. For SLN, this appears to be due to Arg^3^ and Glu^7^, which draws the N-terminus of the transmembrane helix toward the inhibitory groove of SERCA (**Figure 7**). In contrast, the C-terminal end of SLB’s transmembrane helix more closely interacts with SERCA. This appears to be facilitated by a short orthogonal α-helix at the base of SERCA’s inhibitory groove where Pro^952^ of SERCA allows a close approach of Tyr^29^ of SLB. The surrounding residues in SLB (Leu^25^, Trp^26^, & Tyr^29^) and SERCA (Pro^952^, Leu^953^, Pro^954^, & Ile^956^) allow for more efficient packing of SLB in the inhibitory groove, and this may account for the super-inhibitory properties of this peptide. The two most inhibitory SLB peptides, bumble bee and tadpole shrimp, have a tyrosine residue at this position (Tyr^42^ in *Ts*SLB and Tyr^29^ in *Bb*SLB)). The water flea SLB peptide is also super-inhibitory, though slightly less than the other SLB peptides, and it has a cysteine residue at this same position. Compared to the valine found in PLN and SLN (Val^49^ and Val^26^, respectively), the cysteine may also pack more efficiently against the short orthogonal α-helix at the base of SERCA’s inhibitory groove.

## DISCUSSION

The SLBs are a family of peptide orthologs that includes the mammalian regulatory subunits PLN and SLN. While the SLB peptides are conserved from insects to humans, there is a large degree of sequence variation within the family (1). By comparison, PLN and SLN are highly conserved among mammals and they are paralogs with distinct functional and physiological characteristics. This raises the question of whether the SLB peptides possess PLN-or SLN-like functional properties. The goal of this work was to evaluate the breadth of sequence variation in this family of regulatory peptides and how they alter the SERCA calcium pump. This was done by reconstituting three different insect SLB peptides with SERCA into proteoliposomes and measuring changes in ATPase activity of SERCA. Molecular modeling, protein-protein docking, and molecular dynamics simulations were then used to gain novel insight into SERCA inhibition by the SLB peptides.

From these experiments, it was found that the SLB peptides regulated SERCA with two distinct mechanistic features. All three peptides were super-inhibitory, with bumble bee and tadpole shrimp SLB being the most potent SERCA inhibitors. This manifested as a large increase in K_Ca_ reflecting a much lower apparent affinity of SERCA for calcium (**Figures 2 & 3**). We hypothesize that the SLB peptides are normal calcium pump regulators in their native tissues, and that the combination with mammalian SERCA1a accounts for the super-inhibition (see below). In addition to this mechanistic effect on the apparent calcium affinity of SERCA, the SLB peptides also impacted the V_max_ of SERCA. The SLB peptides differed in their effect on SERCA’s V_max_, with bumble bee and tadpole shrimp SLB decreasing the maximal activity of SERCA and water flea SLB increasing the maximal activity of SERCA. Putting these results in context of the functional effects of PLN and SLN on SERCA, the bumble bee and tadpole shrimp SLB peptides exhibited more SLN-like characteristics (decrease in V_max_) and water flea exhibited more PLN-like characteristics (increase in V_max_). As a potential explanation for this observation, the water flea SLB peptide is more oligomeric than the other peptides with discrete trimeric, tetrameric, and pentameric species visible by SDS-PAGE (**Figure 2**). Water flea SLB also has the highest sequence identity to PLN (**Figure 4**).

**Figure 3:**
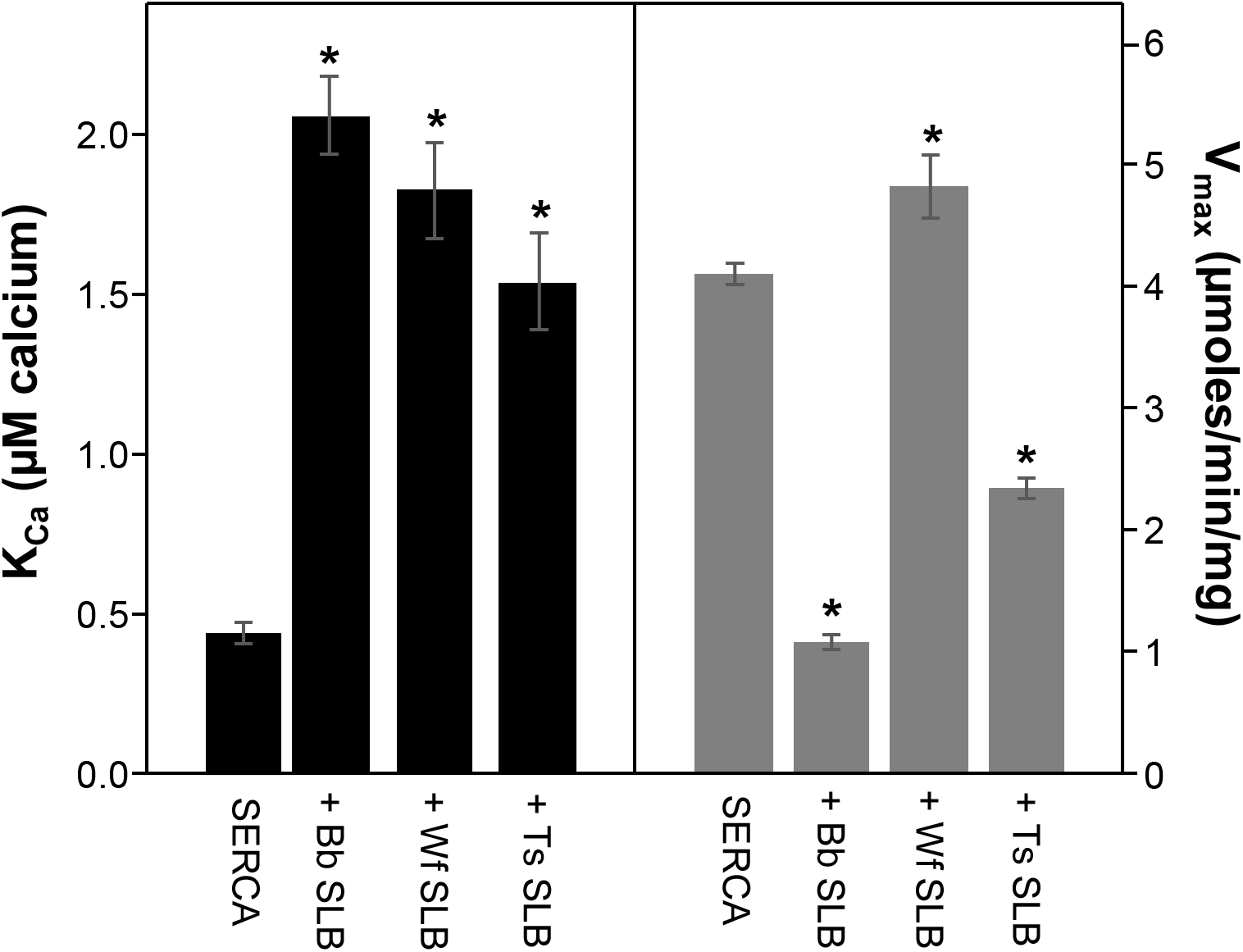
Kinetic parameters determined for SERCA in the absence and presence of SLB peptides. This data is also shown in Table 1. The K_Ca_ values are shown as black bars and the V_max_ values are shown as grey bars. Note that all SLB peptides increase K_Ca_ (decrease SERCA’s apparent affinity for calcium). Bumble bee (Bb) and tadpole shrimp (Ts) SLB decrease V_max_, while water flea (Wf) SLB increases V_max_. Comparison of K_Ca_ and V_max_ was carried out using one-way analysis of variance (between subjects), followed by the Holm-Sidak test for pairwise comparisons (* p < 0.001 compared to SERCA alone).

**Figure 4:**
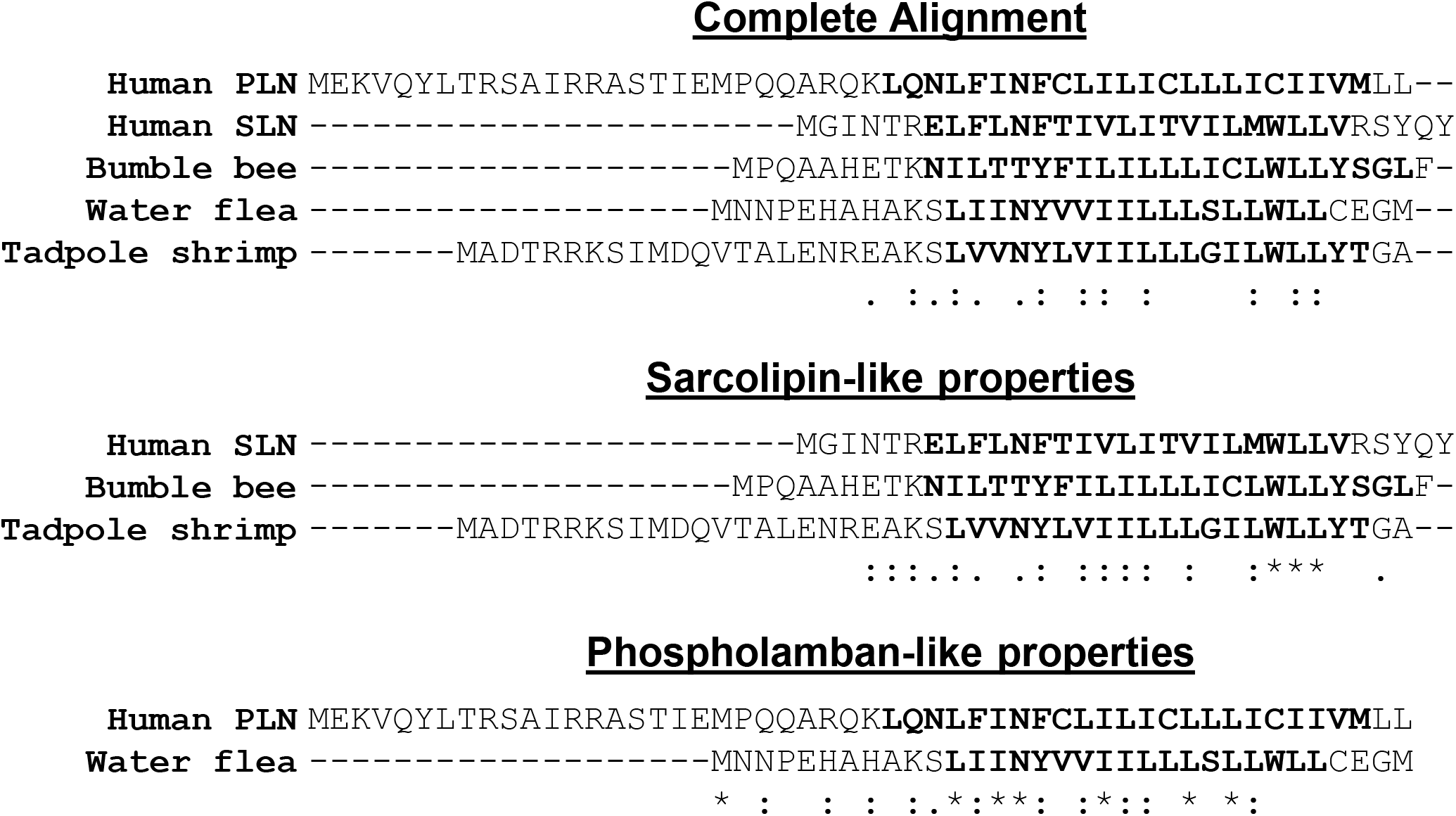
Sequence alignments of the three SLB peptides with PLN and SLN. Shown is an alignment of all five peptides, as well as alignments of the SLN-like SLB peptides with SLN and the PLN-like SLB peptide with PLN.

The SLB peptides used in the present study have similar overall sequences, though they share only 11 identical residues (**Figure 1**). Most notable is the Trp^23^Leu^24^Leu^25^ (WLL) sequence of SLN that is conserved in all three SLB peptides. In the super-inhibitory SLB peptides, this WLL sequence positions a tyrosine residue (Trp^26^Leu^27^Leu^28^ and Tyr^29^ in bumble bee SLB) against the short orthogonal helix at the base of the SERCA inhibitory groove (**Figure 7**). Most notable, Trp^26^ closely approaches the short orthogonal helix in SERCA due to the presence of Pro^952^. This latter residue is Val^952^ in insect calcium pump Ca-P60A, which would be expected to limit the position of Trp^29^ and the penetration of the SLB helix in the inhibitory groove. Thus, Pro^952^ in SERCA likely accounts for the super-inhibition by the SLB peptides, with the additional caveat that the tyrosine residue (Tyr^29^ in bumble bee SLB) packs more efficiently with the short orthogonal helix than the corresponding residues Val^49^ and Val^26^ in PLN and SLN, respectively. While these observations support the effect of the SLB peptides on the apparent calcium affinity of SERCA, the effects of the SLB peptides on the maximal activity of SERCA are consistent with our previous studies (23,24). Both PLN and SLN interact with an accessory site on SERCA (transmembrane segment M3), which appears to account for the effect on the V_max_ of SERCA. SLN interacts as a monomer and decreases V_max_ (24) and PLN interacts as a pentamer and increases V_max_ (23). The PLN-like water flea SLB peptide is more oligomeric than the two SLN-like SLB peptides (**Figure 2**) and it more closely resembles PLN in sequence (**Figure 4**). This explains the ability of water flea SLB to increase the V_max_ of SERCA.

**Figure 5:**
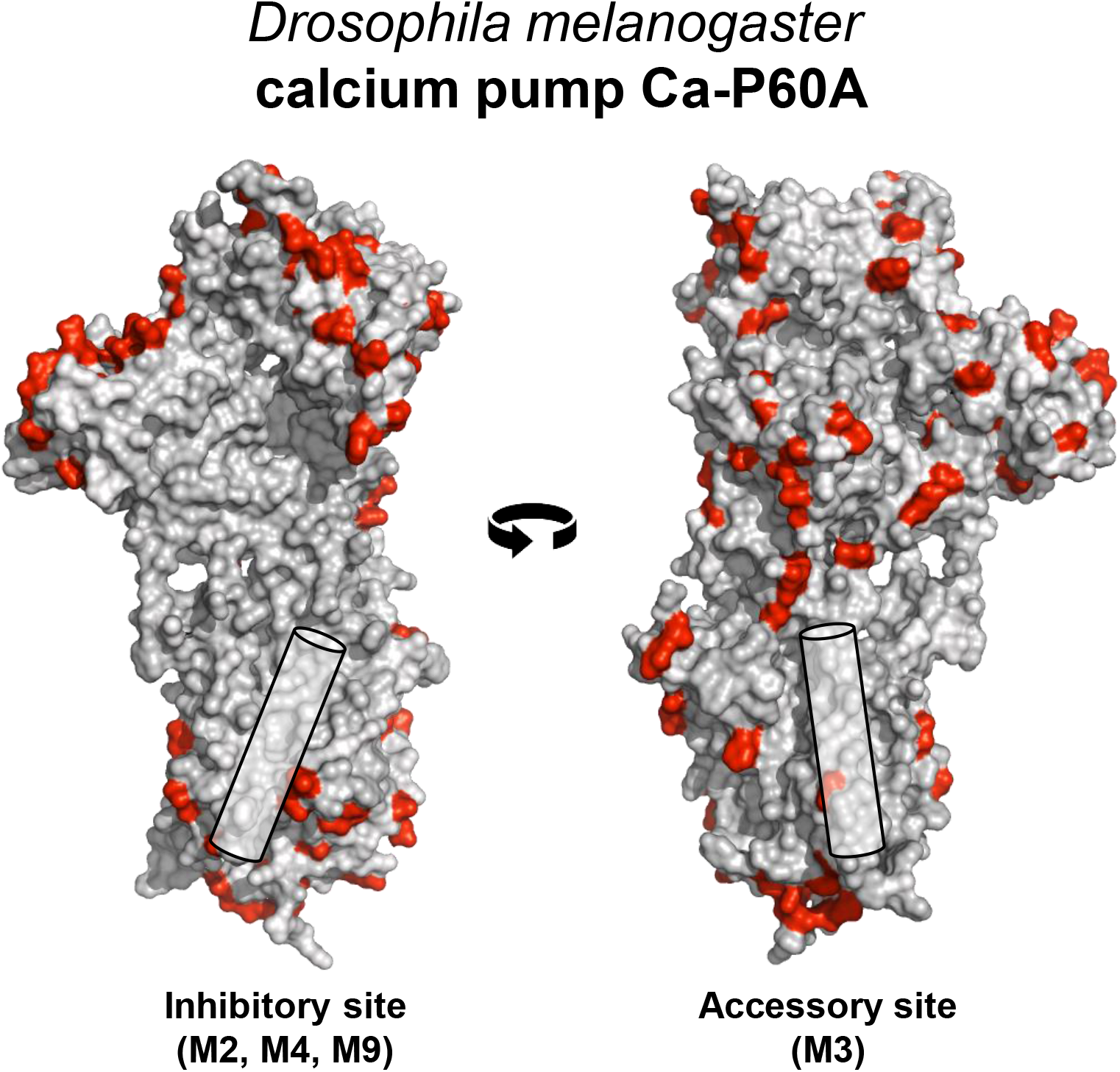
Homology model of the insect calcium pump Ca-P60A. The model was built using Swiss-Model based on the calcium-free E2 state of SERCA (PDB code 3AR4). The model is shown in surface representation with non-conserved residues in red. The inhibitory binding groove is shown on the left and the M3 accessory site on the right. The residues in Ca-P60A that differ from SERCA include Val^952^, Ser^954^, and Thr^955^ at the base of M9 and Ala^273^ on M3.

**Figure 6:**
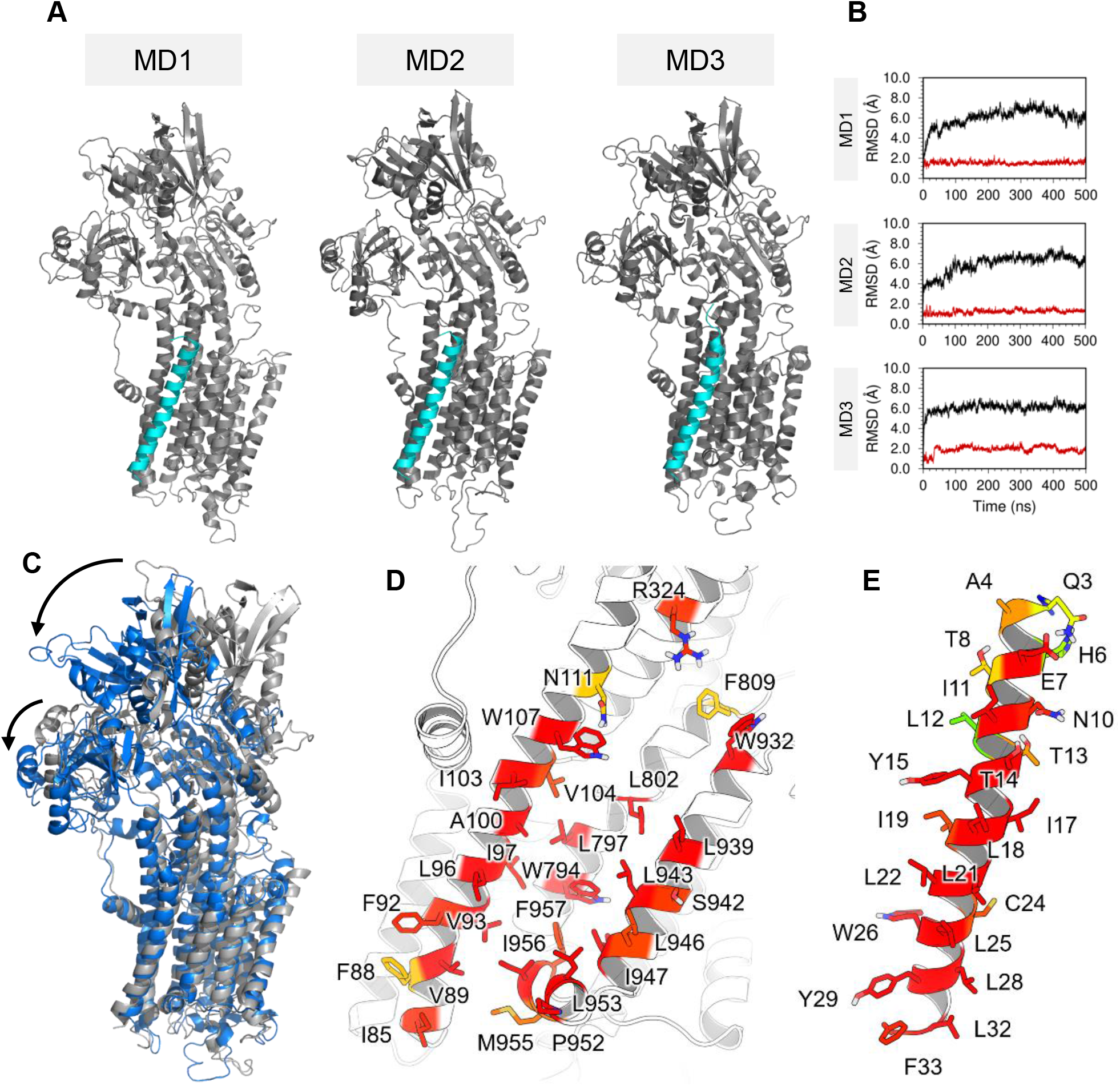
Molecular dynamics simulations of the SERCA-SLB complex. (**A**) Three independent 500 ns simulation replicates of SERCA-SLB complexes (MD1-3). (**B**) Backbone RMSD of SERCA *(black’)* and bumble bee SLB *(red)* for the three simulation replicates. (**C**) Comparison of a representative starting model of the SERCA-SLB complex *(grey cartoon representation)* and a representative model of the final complex after 500 ns simulation for all three replicates *(blue cartoon representation).* Note the movement of the N and A domains of SERCA toward the membrane plane (longer arrow indicates N-domain movement; shorter arrow indicates A-domain movement). (**D**) and (**E**) Residues of SERCA and SLB that form the contact interface with a distance cut off of 3.0 Å.

**Figure 7:**
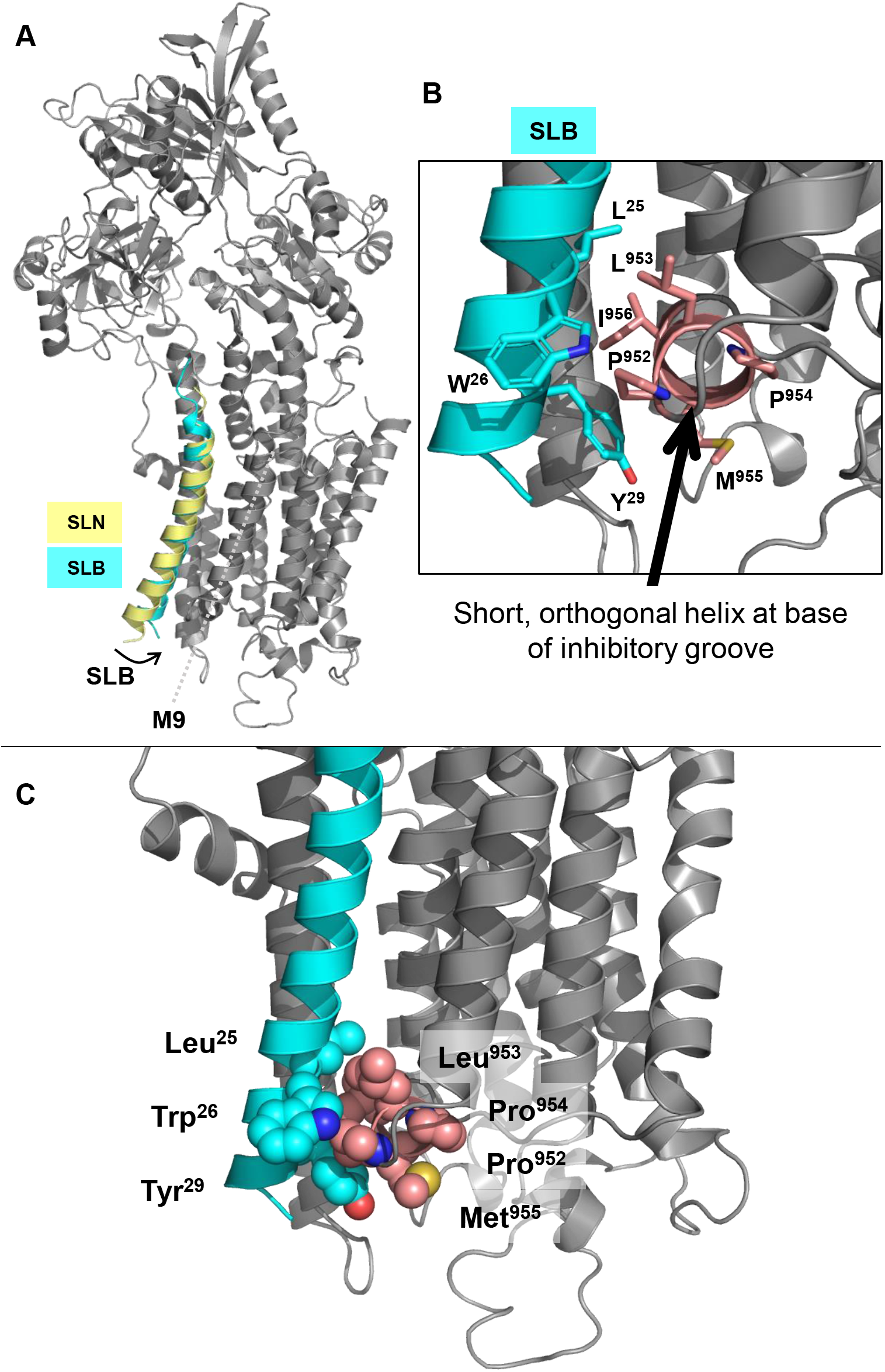
Comparison of SLN and SLB binding to SERCA. (**A**) The SERCA-SLB complex after 500 ns MD simulations is compared to the binding mode of SLN from the known crystal structures. SERCA is shown in grey cartoon representation, SLB is in cyan, and SLN is in yellow. Note the inclination of the C-terminal end of SLB toward the inhibitory grove of SERCA. (**B**) Close up view of the short orthogonal helix at the base on SERCA’s inhibitory groove (pink). Important residues are indicated, particularly Tyr^29^ of SLB, which packs against the short orthogonal helix due to the presence of Pro^952^ of SERCA and a closer approach of Trp^26^ of SLB. (**C**) A similar view as in panel (B) with the residues shown in sphere representation. Note the closer approach of SLB Trp^26^ and Tyr^29^ to SERCA Pro^952^ and the short orthogonal helix.

Despite the breadth of diversity among the ancient family of SLB peptides, they behave like their mammalian counterparts PLN and SLN. We observed that water flea SLB increased the maximal activity and decreased the apparent calcium affinity of SERCA, which is the same effect seen for PLN (14). In contrast, tadpole shrimp and bumble bee SLB decreased both the maximal activity and the apparent calcium affinity of SERCA, which is the same effect seen for SLN (15). When aligning the SLN-like or PLN-like SLB peptide sequences to SLN or PLN, water flea SLB stood out as being most like PLN albeit the overall conservation was limited. All three SLB peptides are super-inhibitors in terms of their effect on the apparent calcium affinity of SERCA. Based on molecular modeling and MD simulations, the SLB peptide has a slightly different inclination in the binding grove of SERCA formed by M2, M6, and M9. The closer approach of the C-terminal end of the transmembrane helix in the region of a conserved WLL motif allows positioning of a tyrosine (bumble bee & tadpole shrimp) or a cysteine (water flea) at the base of SERCA’s inhibitory groove. It is interesting that two SLB peptides elicit a distinctly SLN-like effect on SERCA, though they lack the highly conserved “RSYQY” motif found in mammalian SLN that is so integral to its inhibitory action (15). That said, the SLB peptides are super-inhibitory, so not truly PLN-or SLN-like. This designation is meant to convey the impact of the peptides on the maximal activity of SERCA.

For the PLN-like peptide, water flea SLB, it stands out as the more oligomeric of the SLB peptides. At first glance, tadpole shrimp SLB more closely resembles PLN in length, secondary structure, and charge (**Figure 1**). Secondary structure prediction (26) suggests a short helical region in the N-terminus (residues ~4-15), in addition to the helical transmembrane domain (residues ~24-43), and the overall charge distribution in the cytoplasmic domain is similar (Asp, Glu, Arg, & Lys residues). These features are similar to PLN and consistent with the model for how PLN influences the V_max_ of SERCA (23). However, tadpole shrimp SLB happens to be the least oligomeric peptide with the primary species being a monomer and a small amount of dimer as visualized by SDS-PAGE (**Figure 2**). In contrast, water flea SLB forms an array of oligomeric species with the primary species being a dimer, but monomers, trimers, tetramers, and pentamers are clearly visible. Since the pentameric form of PLN has been found to increase the V_max_ of SERCA (23), this likely underlies the PLN-like effect of the water flea SLB peptide on the V_max_ of SERCA.

The inhibitory groove of SERCA, formed by transmembrane segments M2, M6, and M9, can bind a variety of regulatory peptides including PLN, SLN, and the family of “regulins” (5–7,27). SERCA binding is not very specific as a variety of nonspecific peptides can also inhibit SERCA (28,29) and non-inhibitory mutants of PLN can still bind (30). Moreover, docking and steered molecular dynamics simulations suggest that PLN can bind in several favorable orientations (31) and that the regulatory interaction is not defined by a single discrete state. While this makes rational design of novel SERCA regulators challenging, the present results provide insight into key structural determinants of regulatory function. Potential therapeutic applications include super-inhibitory peptides in the targeted treatment of cancers and non-inhibitory or stimulatory peptides in restoring SERCA function in diseases such as skeletal muscle wasting diseases and heart failure (32). In particular, the use of non-inhibitory mutants of PLN has been proposed as a potential therapeutic approach in diseases such as heart failure where SERCA-dependent calcium cycling is dysregulated (33,34). A variety of synthetic peptides have been shown to inhibit SERCA to varying degrees (28,29). For the naturally occurring peptides, single amino acid substitutions can yield non-inhibitory variants (e.g. PLN Leu^31^-Ala) that still bind to SERCA. However, peptides like PLN that also activate SERCA – i.e. increase the maximal activity – are rare. Besides PLN and some mutant forms of PLN (14,17), no other peptides have been shown to directly activate SERCA. Water flea SLB demonstrates that a distinct peptide can activate SERCA at higher calcium concentrations and it may be possible to design and tailor this peptide for a specific effect on SERCA. Ideally, the peptide would increase the V_max_ of SERCA without altering the apparent calcium affinity (K_Ca_). This would be a potential therapeutic for improving contractility in failing myocardium where the underlying pathology involves elevated cytosolic calcium and reduced pumping efficiency of the heart. Such peptides may be more amenable to gene therapy approaches or as peptide biologics designed to target the cardiovascular system.

A final perspective is the importance of these SLB regulators in invertebrates and their role in calcium signaling. It is known that Ca-P60A is essential for invertebrate calcium homeostasis and that diminished Ca-P60A function leads to temperature sensitivity, decreased heart rate, and altered cardiac rhythm in *D. melanogaster* (35–37). While there is no evidence as to why Ca-P60A needs to be regulated, it likely allows control of contractility, energy metabolism, and thermogenesis in invertebrates. It has also been observed that excess or removal of SLB leads to arrhythmias and altered calcium transients, and that the absence of SLB (and consequently unregulated calcium pump activity) can be rescued by decreasing Ca-P60A activity. Thus, both the Ca-P60A calcium pump and its SLB regulatory subunits need to be present and functional for proper calcium homeostasis and this relationship has persisted for 550 million years of evolution.

## EXPERIMENTAL PROCEDURES

### Materials

All reagents were of the highest purity available including octaethylene glycol monododecyl ether (C_12_E_8_; Nikko Chemicals Co., Ltd, Tokyo, Japan); egg yolk phosphatidylcholine (EYPC), phosphatidylethanolamine (EYPE) and phosphatidic acid (EYPA) (Avanti Polar Lipids, Alabaster, AL); reagents used in the coupled enzyme assay (Sigma-Aldrich, Oakville, ON Canada).

### Expression and purification of recombinant SLB

Recombinant SLB peptides were expressed and purified as previously described (18) with an additional organic extraction step. Following protease digestion of the maltose-binding protein and SLB fusion protein, trichloroacetic acid was added to a final concentration of 6%. This mixture was incubated on ice for 20 minutes. The precipitate was collected by centrifugation at 4°C and subsequently homogenized in a mixture of chloroform-isopropanol-water (4:4:1) and incubated at room temperature for 3 hours. Recombinant SLB was enriched in the organic phase, dried to a thin film under nitrogen gas, and then solubilized in 7 M GdnHCl. Reverse-phase HPLC was performed as described (18).

### Co-reconstitution of SLB and SERCA

Lyophilized SLB peptide (75 μg) was solubilized in a mixture of chloroform-trifluroethanol (2:1) and mixed with lipids (400 μg EYPC, 50 μg EYPE, 50 μg EYPA) from stock chloroform solutions. The peptide-lipid mixture was dried to a thin film under nitrogen gas and desiccated under vacuum overnight. The peptide-lipid mixture was hydrated in water at 37 °C for 30 min, cooled to room temperature, and buffer was added (final concentrations 20 mM imidazole pH 7.0; 100 mM KCl; 0.02% NaN_3_). The peptide mixture was solubilized by addition of C_12_E_8_ detergent (0.2 % final concentration) with vigorous vortexing. Detergent-solubilized SERCA was added (500 μg in a total volume of 200 μl) and the reconstitution was stirred gently at room temperature. Detergent was slowly removed by the addition of SM-2 biobeads (Bio-Rad, Hercules, CA) over a 4-hour time course (final weight ratio of 25 biobeads to 1 detergent). After complete detergent removal, the reconstitution was centrifuged over a 20-50% sucrose step-gradient for 1 h at 100,000g. The resultant layer of reconstituted proteoliposomes at the 20-50% sucrose interface was removed, flash-frozen in liquid-nitrogen, and stored at −80 °C. Quantitative SDS-PAGE was used to determine the SERCA (10% polyacrylamide gels) and SLB peptide (15% polyacrylamide gels) concentrations (38).

### ATPase activity assays of SERCA reconstitutions

ATPase activity of the co-reconstituted proteoliposomes was measured using a coupled-enzyme assay over a range of calcium concentrations from 0.1 μM to 10 μM (14,17). The K_Ca_ (apparent calcium affinity) and V_max_ (maximal activity) were determined by fitting the data to the Hill equation (Sigma Plot software, SPSS Inc., Chicago, IL). Statistical uncertainties were estimated using the standard error of the mean for a minimum of three independent reconstitutions. Comparison of Kea and V_max_ for SERCA in the absence and presence of SLB peptides was carried out using one-way analysis of variance (between subjects), followed by the Holm-Sidak test for pairwise comparisons.

### Molecular modeling of Drosophila melanogaster Ca-P60A

SWISS-MODEL, a fully automated protein structure homology-modeling server (19), was used to generate a molecular model of the *D. melanogaster* calcium pump Ca-P60A (UniProt ID P22700, ATC1_DROME). Ca-P60A served as the target sequence and the calcium-free E2 state of SERCA served as the modeling template (PDB code 3AR4).

### Molecular Dynamics Simulations

A structural model of *B. terrestris* SLB peptide was built with SWISS-MODEL and PEP-FOLD 3.5 (39) servers using SLN structure as a template (PDB code 3W5A). The model was aligned with SLN in the crystallographic structure to generate the complex with SERCA. A total of 23 Cα atoms were aligned computing a 0.815 Å root mean square deviation (RMSD). The spatial arrangement of the SERCA-SLB complex in a lipid membrane was assigned using the Positioning of Proteins in Membrane (PPM) web server (40). The model was inserted in a 1-palmitoyl-2-oleoyl-sn-glycero-3-phosphocholine (POPC) lipid bilayer containing a total 420 molecules using the membrane builder module of CHARMM-GUI web server (41,42). The systems were solvated using a TIP3P water model with a minimum margin of 20 Å between the protein and the edges of the periodic box in the z-axis. Potassium and sodium ions were added to reach a concentration of 150 mM and neutralize the total charge of the system.

Molecular dynamics (MD) simulations of SERCA-SLB were carried out by using the Amber ff14SB (43) and Lipid 17 force field topologies and parameters implemented in Amber 18 and AmberTools package (44). The systems were energy minimized and equilibrated following the six-step preparation protocol recommended by CHARMM-GUI (45). The temperature was maintained at 310 K with Langevin thermostat algorithm and the pressure was set to 1.0 bar using the Monte Carlo barostat. All bonds involving hydrogens were constrained using the SHAKE algorithm. Three independent 500 ns MD replicates of the SERCA-SLB complex were performed. For the MD analysis, AMBER MD trajectories and coordinates were converted to GROMACS files using MDAnalysis python library (46). We calculated the backbone RMSD of SERCA and SLB for the whole trajectories. The frames of the last 300 ns of each simulation were merged and submitted to a backbone RMSD clustering analysis using a cutoff value of 2.5 Å to retrieve the most representative structure of the complex. The independent and merged MD trajectories were further used to calculate the occupancy fraction of SERCA residues located within 3.0 Å of SLB using the select built-in tool implemented in GROMACS (47).

## DATA AVIALABILITY

All data are contained within the manuscript.

## FUNDING ACKNOWLEDGEMENTS

This work was supported by a grant from the National Institutes of Health (R01HL092321) to HSY and SLR. JJB was supported by a 75^th^ Anniversary Award from the Faculty of Medicine & Dentistry, University of Alberta. LME-F was supported by a grant from the National Institutes of Health (R01GM120142). MJL was supported by grants from the Canadian Institutes of Health Research and the Natural Sciences and Engineering Research Council of Canada.

